# The diffusion of cooperative and solo bubble net feeding in Canadian Pacific humpback whales

**DOI:** 10.1101/2024.12.12.628166

**Authors:** Janie Wray, Éadin N. O’Mahony, Grace Baer, Nicole Robinson, Archie Dundas, Oscar E. Gaggiotti, Luke Rendell, Eric M. Keen

**Author notes:** Janie Wray and Éadin O’Mahony share first authorship. **AUTHORS’ CONTRIBUTIONS JW:** conceptualization, data curation, funding acquisition, project administration, writing– review & editing; **ÉOM:** conceptualization, data curation, funding acquisition, formal analysis, investigation, methodology, project administration, validation, visualization, writing– original draft, writing– review & editing; **GB:** data curation, writing– review & editing; **NR:** data curation; **AD:** data curation; **OEG:** investigation, supervision, writing– review & editing; **LR:** conceptualization, investigation, methodology, supervision, writing– review & editing; **EMK:** conceptualization, data curation, investigation, visualization, writing– original draft, writing– review & editing. **DATA ACCESSIBILITY** Code and data associated with this manuscript are available at: https://github.com/eadinomahony/humpback-bubble-netting-NBDA. **ETHICS STATEMENT** This work was carried out under the Department of Fisheries and Oceans research permit DFO XR 83 2014.

## Abstract

Cooperative foraging, a mutualistic resource acquisition behaviour observed across diverse taxa, can drive social network structure, as it typically involves high rates of interactions between individuals. Understanding the mechanisms and distribution of such behaviours is important to elucidate the drivers of social organisation. Bubble net feeding (‘bubble netting’) is a specialised foraging technique practised by certain humpback whale (*Megaptera novaeangliae*) populations globally. Over 20 years of study in the northern Canadian Pacific, we observed the diffusion of two morphs of this behaviour: social cooperative and independent, or ‘solo’, bubble netting. Network-based diffusion analyses – a tool developed to test for social learning – find strong evidence for the social learning of bubble netting when a static social network is used, even after accounting for individual-level traits such as site fidelity and sex (10.6 ×10^3^ to 35.4×10^3^ times more support for social versus asocial learning; p < 0.0001). However, we could not completely exclude homophily due to the inherent sociality of this cooperative foraging behaviour. Nonetheless, it is clear that the rapid diffusion of bubble netting throughout this population has significant consequences for its social structure, which should be information considered by those planning the conservation of this threatened population.

## INTRODUCTION

Animal culture, in which information and behaviours are acquired and shared through social learning [1–5], is a form of biodiversity with both intrinsic value and practical importance [1,5–8]. Learned behaviours augment population viability, bolster individual survival, reinforce group cohesion, increase intraspecific diversity, facilitate ecological specialisation, enhance resource utilisation, and promote resilience to ecosystem change [1,3,6–13]. Cultural traditions, such as learned migratory routes, breeding habitat, and feeding areas, also determine species’ responsiveness – or lack thereof – to habitat alterations within their range [1]. For these reasons, the importance of animal culture and social learning to biological conservation is receiving increasingly widespread recognition [1,8,14–16].

The social learning of foraging techniques can be particularly instrumental to a species’ survival and ecological resilience [13]. Novel techniques can improve rates of prey capture or provide access to new types of prey, thus expanding viable habitat while reducing vulnerability to shifts in prey supply and distribution [13,17]. The diffusion of new feeding strategies through social groups has been described for various taxa, from fish [18,19] to birds [20–22] and mammals [13]. Mammalian examples of socially learned feeding techniques are particularly common among primates [23–29] and cetaceans [7,11,30–33], particularly the odontocetes [9,34–37].

Social learning, the mechanism therefore underpinning cultural evolution, has been typically categorised into the occurrence of learning as 1) a direct result of *observations* of a behaviour; or 2) a consequence of ‘*interactions with another animal* (typically a conspecific) or its products*’* (emphasis added; [3]). Given a vast range of cooperative behaviours across taxonomic groups, documented particularly well in cetaceans [38–44], we emphasise social learning as a result of interacting with a conspecific. Social learning can be seen as a highly dynamic process of observation and simultaneous interaction with the behaviour of ‘informed’ individuals and the products of their behaviour, i.e. a process of ‘learning by doing’.

With few exceptions (i.e., migratory routes of southern right whales, *Eubalaena australis* [45], and possibly tread-water feeding in Bryde’s whales, *Balaenoptera edeni* [46]), the humpback whale, *Megaptera novaeangliae*, is the only baleen whale for which there is currently good evidence of widespread social learning. While the cultural evolution of humpback song is perhaps the most widely known [47,48], humpback whales also innovate and socially transmit feeding techniques, such as lobtail feeding in the North Atlantic [30,49] and trap feeding near Vancouver Island, Canada [13]. Other feeding strategies of this generalist predator include lunge-feeding [50,51], flick-feeding [50,52], bottom-feeding [53,54], and bubble net feeding (hereafter ‘bubble netting’) [44,50,55–60]. It is possible that some of these other techniques are also socially learned, but to date have not been demonstrated as such.

Bubble netting is a well described foraging strategy performed by humpback whales across different feeding grounds and ocean basins. The behaviour itself is a multi-step task: diving beneath a shoal of schooling fish or krill and emitting a stream of bubbles through the blowholes whilst swimming in a spiral or circle to spin a “net” or curtain of bubbles, entrapping the prey [61]. Sometimes this is accompanied by specific vocalisations produced at a frequency shown to vibrate the swim bladders of the target prey; or act as sonic traps in tandem with the bubble net, where a ‘wall of sound’ is formed with a relatively quieter interior at the centre of the bubble net [62]. The feeding bout is concluded with a vertical or horizontal lunge through the prey patch at or near the surface of the water. This behaviour is often performed by individual whales [57–59], but it can also be performed by multiple individuals coordinating their behaviour with role differentiation and cooperation [44,50,56,57,60]. While the relative benefits of bubble netting individually compared to cooperatively are not well understood, Mastick *et al*. suggest that in larger bubble net groups, each individual has decreased exertion, ultimately expending less energy in relation to amounts of prey captured [57]. An important point is that group bubble netting is genuinely cooperative, with division of labour, not simply a close spatial aggregation of individually feeding whales.

Given that cooperation usually involves coordinated actions and at times a division of labour and further role specialisation [43,63], stable social associations and increased group size might then lead to a higher net benefit in cooperative foraging [43]. It has been suggested that groups of bubble netting humpback whales have role specialisation across foraging bouts (where a ‘foraging bout’ is considered to be one iteration of the casting of a bubble net with subsequent lunge feeding by all group members) [57]. Conceivable roles include the production of the bubble net itself, the herding of shoaling fish into the net, the vocalisation of characteristic feeding calls and the ultimate vertical or horizontal lunge feed [44]. However under the definition of a ‘team’ – a cooperative behaviour with a division of labour, requiring individuals to perform different subtasks for the success of the behaviour [63] – bubble netting humpbacks in certain regions show high levels of behavioural plasticity where teamwork is apparent but not strictly necessary for the ‘successful completion’ of the task. For example, the humpback whales foraging in the Kitimat Fjord System (KFS) of northern British Columbia (BC), Canada, show large variability in bubble netting group size and certain individuals switch between cooperative group bubble netting and individual, or ‘solo’, bubble netting.

Here we hypothesise bubble netting to be a behavioural trait underpinning the structure of the social network of an annually resident population of humpback whales [64], with the inherent sociality and cooperation of this behaviour mediating its cultural transmission. The behavioural plasticity of this locally expanding population may contribute to its ability to recover from a history of commercial whaling [60,65], which resulted in the catch of an estimated ∼29,000 humpback whales across the North Pacific in the 20th century [66]. Understanding the mechanisms of resilience of local populations, in light of increasing anthropogenic stressors, is particularly important given more recent evidence of a decline in the North Pacific-wide population [67]. We tested for evidence of social learning of bubble netting using a network-based diffusion analysis (NBDA; [68]) on a dataset of humpback sightings from 2004 to 2023.

## METHODS

### (a) Data collection

Humpback whale photo-identifications and behavioural observations were collected from 2004 to 2023 (n=20 years) within the marine territories of the Gitga’at, Kitasoo/Xai’xais and Haisla First Nations, in the KFS (N 52.8 – 53.5, W 129.6 – 128.5; Fig. 1), BC, Canada. This work was carried out in close collaboration with the Ocean and Lands Department of the Gitga’at First Nation and with the explicit permission of each aforementioned Nation. Whales were observed from shore-based and vessel-based platforms. The vessel-based survey methods we used are detailed in [65] and [56]. Briefly, pre-planned survey routes were conducted between 2004 and 2023 using a 7 m skiff as weather permitted from April to November. When humpback whales were detected, groups were approached with caution, all individuals were counted, location and behaviour noted, and identification photographs of the underside of their tail flukes were collected with standard DSLR cameras and telephoto lenses, following established protocols for this species (e.g., [69]). Fluke photograph capture was non-systematic in order to maximise the number of whales identified.

**Figure 1.**
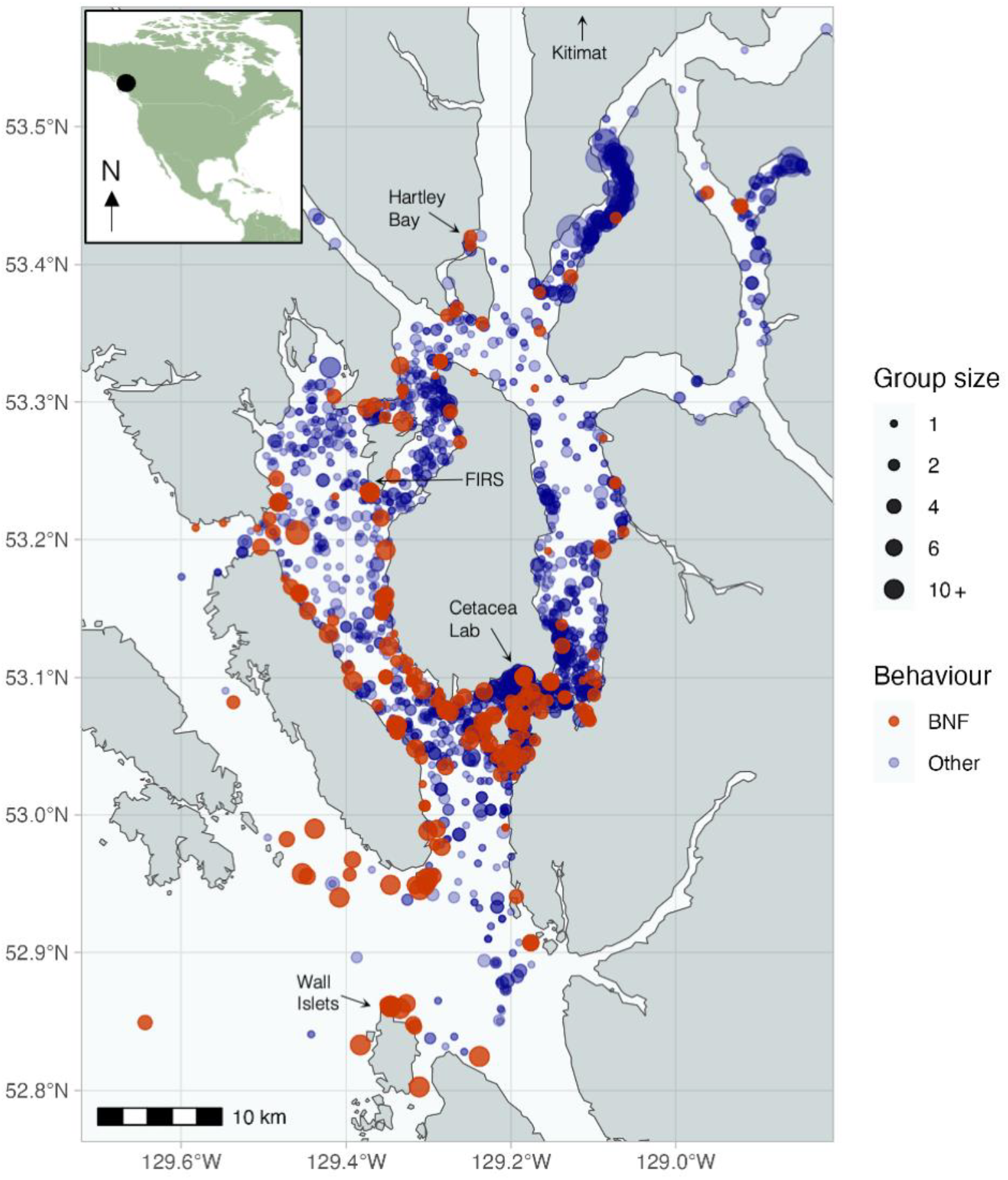
Humpback whale sightings, colour-coded and sized by behavioural state (orange represents sightings of bubble netting, while blue represents sightings of any other behavioural state), in the Kitimat Fjord System, British Columbia, Canada, within the marine territories of the Gitga’at, Kitasoo/Xai’xais and Haisla First Nations. Field stations and population centres are labelled.

Shore-based observations occurred at Cetacea Lab (53° 6’17.31”N, 129°11’37.72”W) from 2014 to 2016, the Wall Islets (52°51’29.10”N, 129°20’28.53” W) from 2014 to 2016, and Fin Island Research Station (FIRS, N 53°13’18.94”N, 129°22’34.77”W) from 2018 to 2023 (Fig. 1). At these stations, observers carried out systematic scans of the aquatic viewshed at regular intervals from sunrise to sunset from May to October. Groups were noted if they came within 500 m of shore and identification photographs were taken and catalogued.

### (b) Data analysis

For analysis, we defined a *sampling period* as a single calendar day of photo-identification effort, which usually consisted of up to 16 hours of active monitoring at shore-based stations and 2 – 10 hours of surveying during vessel-based effort. An *encounter* is defined as a unique observation of a unique group (i.e. if a group of the same social composition was observed twice in one day, as determined through photo-identification, it is only included once; if certain whales from a previous group are seen with other whales in a separate group later in the same day, both encounters are included). A *group* is defined here as individuals that come within two body lengths of each other and coordinate their swimming, diving, and/or ventilation behaviour for at least one surfacing [56,70]. We use the term *population* here to refer to the local population of humpback whales documented within the KFS, unless otherwise specified as referring to the entire North Pacific population. Bubble netting behaviour was easily identifiable due to its many indications at the surface (bubble rings, vertical lunges and often conspicuous group sizes). Females were identified when they arrived with a calf and additional sex information was obtained from blow samples collected using an unoccupied aerial system [71]. All identification photographs were scored for quality and only high-quality photographs were used for annual catalogues of identified individuals [65]. Dyadic association weights were determined with the Simple Ratio Index (SRI, [72]) based on recommendations in [73,74]. Social networks were visualised using all individuals observed on at least 5 occasions using the R package ‘igraph’ [75].

We used a network-based diffusion analysis (NBDA; [68,76]) to test for evidence of cultural transmission of bubble netting in whales encountered 5 or more times (both group and solo events included). The underlying premise of NBDA is that learned behaviours tend to spread more quickly among individuals who spend more time together, and that heightened rates of trait performance will increase the likelihood of behavioural transmission amongst associated conspecifics [68,77]. NBDA models the rate at which ‘naïve’ whales (i.e., those never before observed to perform the behaviour) are first observed bubble netting as a function of their associations with ‘informed’ individuals. The likelihood of the model’s fit is then compared to a null model of asocial learning (i.e., independent discovery or invention), where the social transmission parameter is fixed at zero [76].

We built models using a multiplicative order-of-acquisition diffusion analysis (OADA), as we do not know the exact times of acquisition of the target behaviour and therefore assign a ranked bubble netting acquisition order onto whales in the order in which we documented them performing the behaviour [76]. Newly informed individuals from the same detection window were assigned the same ranked order, i.e. they were tied to each other. All individuals observed bubble netting in the study’s first year during their first sighting (*n* = 12, representing 7% of the individuals seen five or more times and known to perform the behaviour at the end of the study) were assigned a rank as ‘demonstrator’, assumed to seed the population with the behaviour, and therefore were removed from the order of acquisition. We chose three individual level variables (ILVs) to include in the models. All models include the total number of sightings of that individual, standardised to a mean of zero and standard deviation of one, in order to account for sampling bias in our dataset. Models were then built using a combination of site fidelity, as measured by the Standardised Site Fidelity Index (SSFI, range 0-1) [78], and sex, to explore the impacts of these ILVs on estimated social transmission. We chose to use multiplicative models because of our underlying assumption that each ILV would affect social and asocial learning rates equally (e.g. there is no biological evidence we know of to suggest that one biological sex might be better at learning socially or, alternatively at asocial innovation, than the other). Models were compared using ΔAIC, and the lowest ranking model used to further test for the effect of homophily.

To determine the influence of each ILV on social learning, we calculated the summed Akaike weights (Σwi) across all models, the model-averaged parameter estimate with 95% confidence intervals calculated according to Eq. 6.12 in [79] (see Supplementary Materials for equation). Effect sizes are calculated as e^(β/σ)^ for continuous variables (standardised sightings and site fidelity SSFI), where β is the model-averaged parameter estimate and σ is the standard deviation of the unstandardised data. Effect size is calculated as e^(2β)^ for the categorical variable sex where sex is 1 for male, -1 for female and 0 as unknown.

#### Homophily & immigration

Given the inherent sociality of the cooperative bubble netting behaviour, we tested for the putative bias that post-acquisition social associations might have on our ability to test for social transmission. A standard OADA-type model uses one overarching static social network, composed of dyadic social association strengths (in our case SRI values) spanning the length of the study period. However, as this approach does not differentiate between encounters before and after the acquisition events in calculating the social network, it does not distinguish homophily - where animals that share a behaviour tend to be around each other more, perhaps simply because they target the same resource or cooperate in the targeting of the shared resource. To do this we needed to target individuals that (a) acquired the behaviour during the study and (b) had a reasonable estimate available of their pre-acquisition social network. Therefore, to test for the effect of homophily in this context, we split the sightings data into two time periods (2004-2013 and 2014-2023) and calculated a new social network (a matrix of dyadic SRI scores) at each bubble net acquisition event that occurred in the second half of the study, where the individual had been seen at least 5 times before being observed bubble netting. Each iterative social network therefore included the full first half of the study period plus the additional time to acquisition. This dynamic social network was then incorporated into a final OADA model with the format of the most parsimonious static network (i.e. the lowest AICc scoring model of ILV combinations). We included as seeded demonstrators all whales seen bubble netting in the first half of the study period as well as any individuals seen bubble netting on their first sighting in the second half of the study period. Ties were treated as before, with whales observed bubble netting for the first time together being given the same rank in the order of acquisition (i.e. they are assumed to have acquired the trait at the same time). All analyses were conducted in R v. 4.3.1 [80] using custom scripts and R package ‘NBDA’ [77].

## RESULTS

A total of 7,485 photo-identifications were collected during 4,053 encounters across 20 years of survey effort (1,356 days of sampling, 2004 - 2023). Of the 526 identified individual whales, 250 were encountered on ≥5 occasions. 58% of whales were encountered across multiple years (32% in 5 or more years, 15% in 10 or more years). Bubble netting was observed in 254 individuals (48% of the identified population, hereafter referred to as ‘bubble netters’) on 635 occasions. The form of bubble netting observed was similar to that described in southeast Alaska [50,55], in which one individual in the group (or in rare cases more than one; pers. obvs.) produces a distinct tonal moan (400 – 600 Hz) that has come to be known as a ‘feeding call’ [50]. Solo bubble netting does not always include the production of a feeding call (ÉOM, EMK, GB, JW, pers. obvs.). Pacific herring was the only prey observed during cooperative feeding events or indicated from the collection of prey remains (JW and GB pers. obvs.).

Of all 254 bubble netters, 32% (n=82) were bubble netting during our first encounter of them. Of these, 65 individuals (25.6% of all bubble netters) were observed cooperatively bubble netting on first encounter. The remaining 17 individuals (6.7% of all bubble netters) were observed solo bubble netting on first encounter. Therefore, most individuals seen bubble netting on first encounter were doing so cooperatively. The large majority of all the bubble netting events observed were cooperative - 92.4% of all 635 bubble netting events documented during the 20-year study period. Of the 17 individuals first seen in a solo bubble netting context, six (35%) were subsequently observed participating in cooperative bubble netting. Given a lesser prevalence of solo bubble netting overall, a total of 167 solo bubble netting events were documented (7.6% of all bubble netting events), these were performed by a comparatively high number of unique individuals, 93 individuals in total. Of the remaining 172 bubble netters, who were first seen in non-feeding contexts, 151 (88%) of these were cooperatively bubble netting when first observed in a bubble netting context.

When data are filtered to whales seen ≥5 times, 179 individuals were seen bubble netting on 614 occasions (72% among whales encountered ≥5 times). The size of bubble netting groups ranged from 1 to 16 (median=3), with 30% of encounters involving groups of ≥5 whales. Of the 179 bubble netters that were seen ≥5 times in the study period, 151 of them were encountered before they were seen bubble netting for the first time (Fig. S1). There was a relatively steady increase over time in the proportion of the NCCS catalogue that were known bubble netters (Fig. 2A).

**Figure 2.**
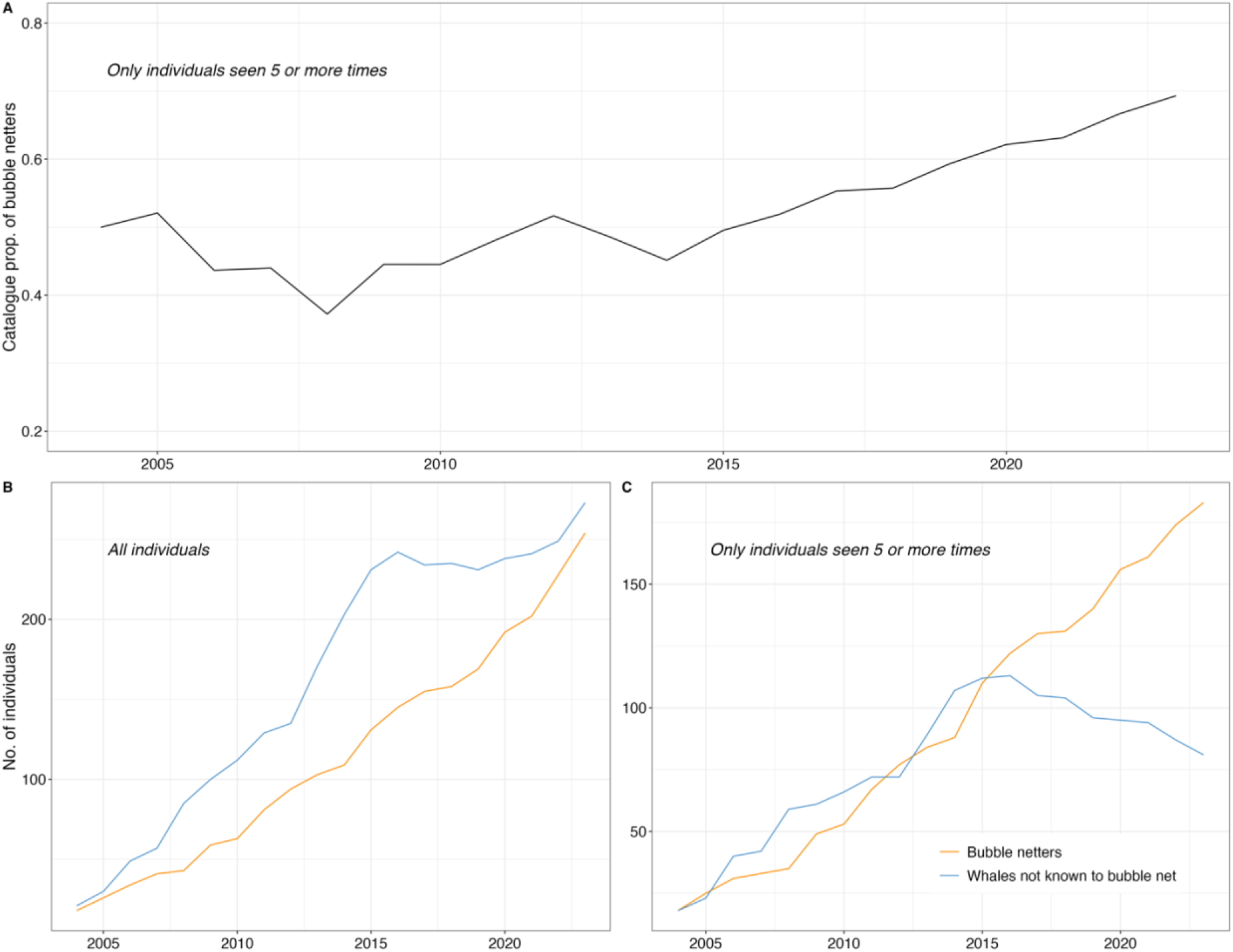
**A)** The proportion of whales who are known to bubble net in the NCCS catalogue each year showing an accelerating spread in the second half of the study period. **B)** The cumulative number of individuals known to the catalogue, seen any number of times (i.e. all individuals), split by whether or not they are known to bubble net (orange = bubble netters; blue = not known to bubble net). **C)** The summative number of individuals known to the catalogue that have been seen five or more times, split as in B).

An important pattern emerges when the set of known individuals in the study is split based on whether or not they have been seen bubble netting up to that point (i.e. becoming informed individuals; Fig. 2B-C). When this data is visualised with all whales, regardless of number of sightings (Fig. 2B), we see a steady increase in both bubble netters (orange) and non-bubble netters (blue), suggesting an immigration of informed individuals, as opposed to social learning upon arrival to the KFS. However, when the less sampled individuals are removed (i.e. those seen less than 5 times in total), the pattern changes: we see a steady increase in both bubble netters and non-bubble netters until 2014, at which point the number of non-bubble netters reverses its trend and begins to decline (Fig. 2C). Given this measure is summative and the behavioural state a binary either/or, the only explanation for this decline is that whales already known to the catalogue are beginning to bubble net having not done so previously. The decline in non-bubble netters is explained by naïve individuals becoming informed and switching into the bubble netter category. This suggests a rapid diffusion of the behaviour throughout the population. This coincides with the severe northeast Pacific Marine Heatwave (2014-2016; [81]), which negatively affected regional humpback whale populations [67,82–84].

Social transmission was strongly favoured in all static NBDA models of bubble netting, with all models of social learning scoring a lower AIC than the equivalent asocial model. The social network used for these models, based on SRI scores, is visualised in Figure 3, with thicker edges depicting social associations significantly more likely than by random chance (p < 0.05). The four social models, differing only in their ILVs, ranged from 10.6×10^3^ to 35.4×10^3^ times more support than their asocial counterparts (likelihood ratio test (LRT) results in Table 1; p < 0.0001). The estimated proportions of acquisition events that result from social transmission (%ST; [77]) vary little across models (min = 55.6%; max = 59.9%) indicating that the model estimates for social learning are not reliant on the inclusion of particular combinations of ILVs. To interpret the effect of each ILV on social learning, we calculated the model-average parameter estimates using the summed Akaike weights (Σw_i_; Table 2). For every additional 18 sightings of an individual (the median number of sightings in the study), there was a 46.8% increase in apparent learning rate, so it was important to try and account for that in all models. There was an 18% reduction in apparent learning rate associated with being male. Compared to an animal with zero site fidelity, an animal with complete fidelity has a 15.7% greater apparent learning rate.

**Table 1.** Results of multiplicative Order of Acquisition Diffusion Analysis (OADA), testing for evidence of social learning in the diffusion of bubble netting throughout humpback whales of the Kitimat Fjord System, British Columbia, Canada. All models treat whales observed bubble netting in year one of the study (2004) to have already acquired the trait before the study began. All models include the standardised number of sightings as an individual level variable (ILV), in order to correct for potential sampling bias. All models assume ILVs to affect the social and asocial learning rates to be equal (multiplicative modelling; see text for details).

| Static Models | AICc | ΔAIC | $w_i$ | MLE (ST) | 95% C.I. | % ST<sup>x</sup> | LRT<sup>†</sup> | p-value<sup>†</sup> |
| --- | --- | --- | --- | --- | --- | --- | --- | --- |
| OADA with std. sightings | 1561.28 | 0.00 | 0.36 | 4.01 | 1.42-10.66 | 55.6% | 20.60 | *** |
| OADA with std. sightings & sex | 1561.32 | 0.04 | 0.36 | 4.70 | 1.71-12.53 | 58.8% | 22.64 | *** |
| OADA with std. sightings, SSFI & sex | 1562.99 | 1.71 | 0.15 | 4.98 | 1.79-13.56 | 59.9% | 23.05 | *** |
| OADA with std. sightings & SSFI | 1563.19 | 1.90 | 0.14 | 4.13 | 1.45-11.11 | 56.2% | 20.76 | *** |

<sup>x</sup> %ST = Percentage social transmission
<sup>†</sup> LRT = Likelihood ratio test for social transmission and associated p-value.

**Table 2.**
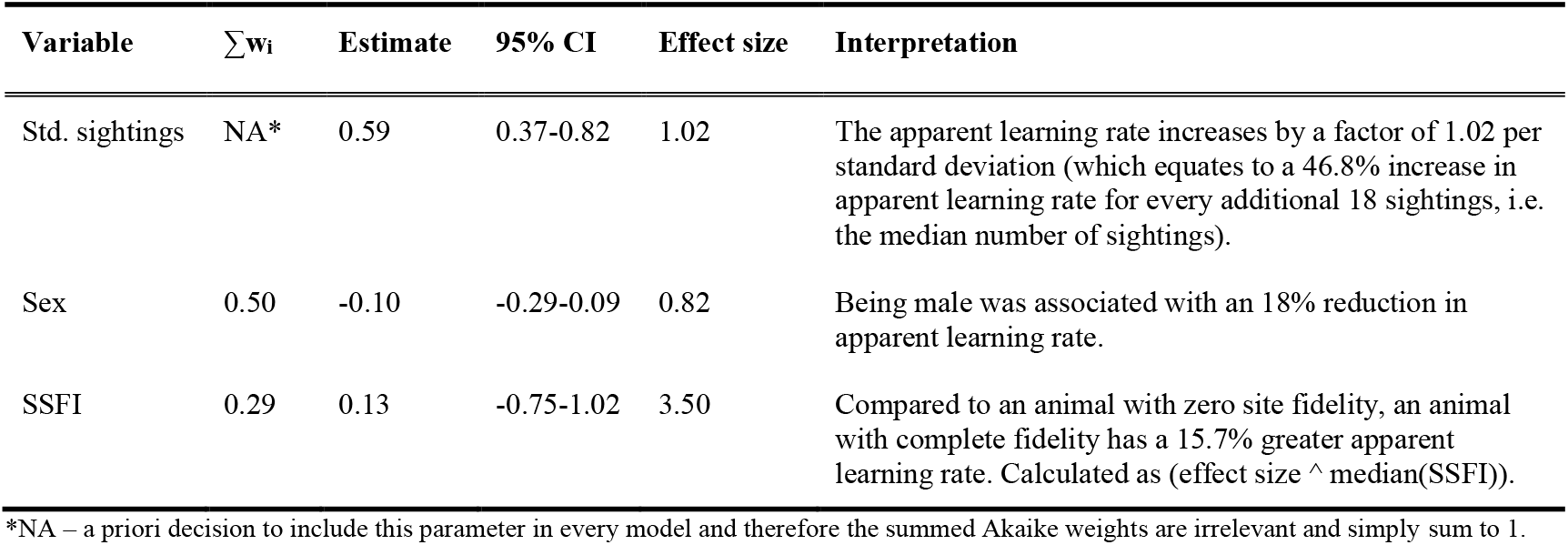
Individual level variables summed Akaike weights (Σw_i_) across all models, the model-averaged parameter estimate with 95% confidence intervals calculated according to Eq. 6.12 in [79]. Effect sizes are calculated as e^(β/σ)^ for continuous variables (sightings and site fidelity SSFI), where β is the model-averaged parameter estimate and σ is the standard deviation of the unstandardised data. Effect size is calculated as e^(2β)^ for the categorical variable sex where sex is 1 for male, -1 for female and 0 as unknown.

| Variable | $\sum w_i$ | Estimate | 95% CI | Effect size | Interpretation |
| --- | --- | --- | --- | --- | --- |
| Std. sightings | NA* | 0.59 | 0.37-0.82 | 1.02 | The apparent learning rate increases by a factor of 1.02 per standard deviation (which equates to a 46.8% increase in apparent learning rate for every additional 18 sightings, i.e. the median number of sightings). |
| Sex | 0.50 | -0.10 | -0.29-0.09 | 0.82 | Being male was associated with an 18% reduction in apparent learning rate. |
| SSFI | 0.29 | 0.13 | -0.75-1.02 | 3.50 | Compared to an animal with zero site fidelity, an animal with complete fidelity has a 15.7% greater apparent learning rate. Calculated as $(\text{effect size} \wedge \text{median}(\text{SSFI}))$. |

\*NA – a priori decision to include this parameter in every model and therefore the summed Akaike weights are irrelevant and simply sum to 1.

**Figure 3.**
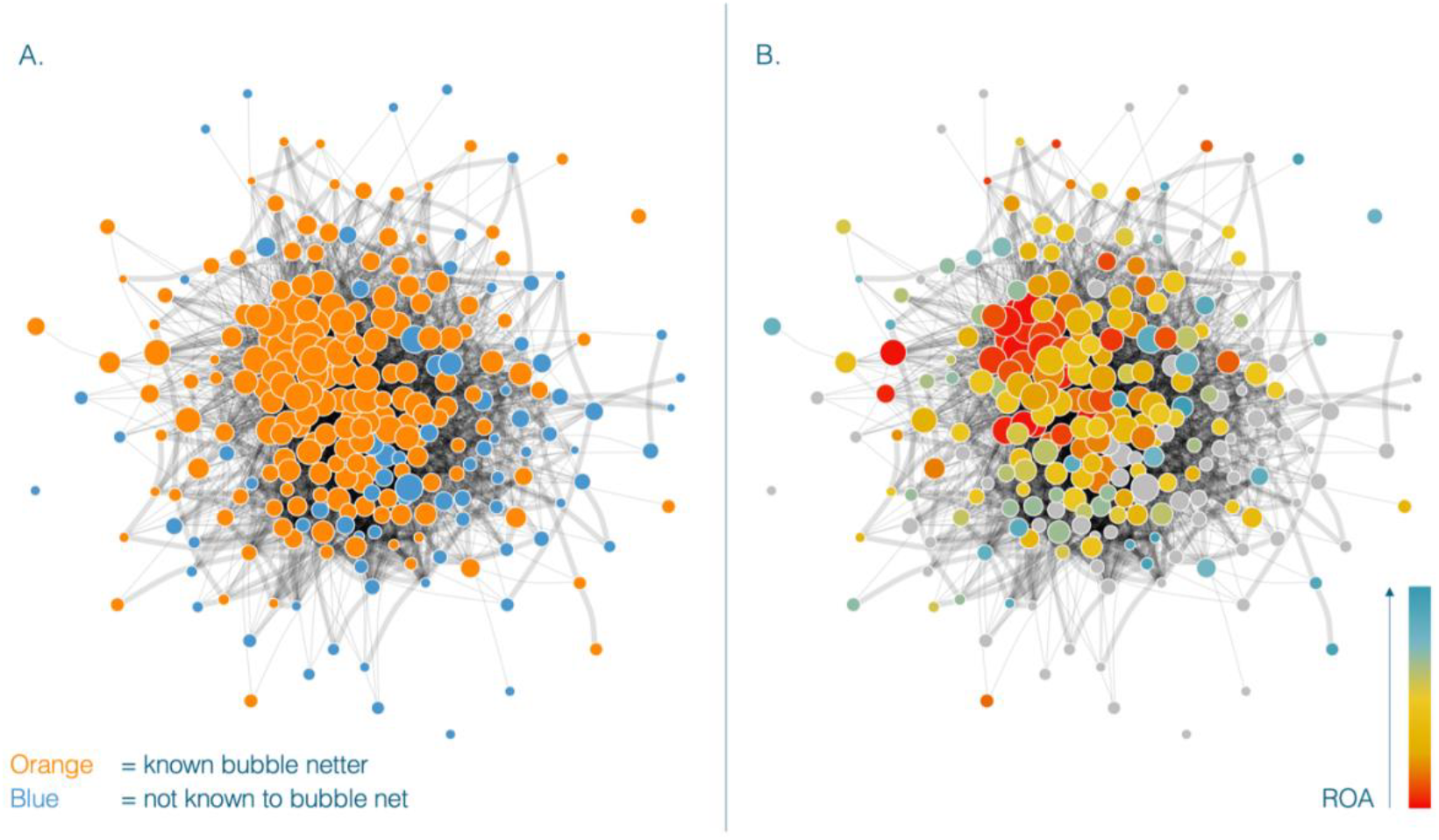
Humpback whale network of social associations (≥5 encounters), weighted by the Simple Ratio Index where thicker edges are social associations significantly more likely than by random chance (p < 0.05). The networks are colour-coded by **(A)** known bubble netting activity (orange nodes are known bubble netters, while blue nodes are not known to bubble net), and **(B)** observed ranked order of acquisition (ROA) of the bubble netting strategy that we observed in humpback whales in the Kitimat Fjord System (warm colours are early acquisition events, moving to the cooler colours which occurred later in the study period). Grey nodes represent whales not known to bubble net. Nodes are scaled by the total number of sightings for each individual, with more sightings increasing the size of the node, and clustered according to the magnitude of their social association measures.

However, the dynamic OADA, which was built to test for the effect of homophily, showed inconclusive results: the asocial model estimated 30.1% of acquisition events occurred by social transmission (%ST). A likelihood ratio test showed that including a social parameter did not significantly improve model fit over the asocial version (LRT = 1.47, p = 0.22). It is important to note that this homophily test had a restricted dataset of 51 individuals acquiring the behaviour, representing 30.5% of the 167 acquisition events of the full dataset, reducing its overall statistical power.

## DISCUSSION

We aimed to test for the role of social transmission in the observed spread of a foraging behaviour in humpback whales. A NBDA of the 20-year dataset showed strong support for social transmission. However, when data were restricted to test for the effect of homophily, the analysis supported asocial and social learning equally. It is difficult to determine whether this is due to a lack of power from a restricted dataset or whether bubble netting is really learned individually followed by a strong homophily effect in the context of cooperation. A third possibility is a confounding in the NBDA because the behaviour, cooperative bubble netting, is itself social. This latter fact however, along with tracking of the dynamics of the behavioural spread, and the overwhelming prevalence of cooperative over solo bubble netting in our data, suggest significant social components. Irrespective of the precise learning trajectories, our analysis clearly shows that the diffusion of bubble netting is very strongly linked with the social structure of this expanding population and is potentially ecologically significant insofar as it affects the distribution and ecological impact of humpbacks foraging in the KFS. We expect the rapid diffusion of primarily cooperative bubble netting to have cascading effects on ecological niche partitioning, and beyond this, a likely effect on the population’s ability to recover to pre-whaling numbers.

Our study took place at the end of a period of population expansion (the North Pacific has since experienced a decline [67]) so it is plausible that population movements are a factor in the spread of this behaviour, rather than learning. However, using almost 4,000 identification photographs, Calambokidis *et al*. found little interchange between different feeding grounds of humpback whales in the North Pacific, and we have found high rates of return and residency amongst the whales foraging in the KFS (Figure S1; [85]). This suggests informed individuals are not simply immigrating to the study region and thereby ‘importing’ the feeding behaviour from elsewhere and instead supports the hypothesis that some whales are learning the behaviour onsite. There is an increasing proportion of bubble netters in our research catalogue (Figure 2A), and a clear switching of better-sampled individuals from a naïve to an informed status in regard to bubble netting ability (Figure 2C). Whales infrequently seen (less than 5 times) masked this trend of behavioural diffusion, so it is important to account for this in network analyses (Figure 2B).

### Learning to cooperate: The crux of distinguishing social learning from homophily

We must be cautious with the interpretation of the results from the NBDA, primarily due to the confounding results between the primary NBDA models showing very strong support for social learning (between 10,000 and 35,000 times more support for the social versus asocial models; p < 0.0001) compared with the dynamic social network used to test for the bias of homophily, which is inconclusive (LRT = 1.47, p = 0.22). It might be premature to conclude that bubble netting is in fact socially learned, despite it being inherently social in most instances (92% of bubble netting events occurred in groups of two or more whales). The results of the homophily test may show that, in fact, homophily is reflected in the social network, such that we find very strong support for social learning when not correcting for its effects. However, this homophily test is vulnerable to failing *because* bubble netting occurs socially, and so we would expect bubble netters to bias their social associations towards other bubble netters (i.e. homophily) in order to feed cooperatively. This suggests that a NBDA might in this context, where the behaviour relies on inherent sociality and cooperation, lack power to detect the signals of social learning.

A literature review of applications of NBDA^1^ demonstrates that, to our knowledge, all behaviours tested with this tool which find support for social learning, are learned socially but performed individually. In cetaceans, examples include lobtail feeding in humpback whales [30]; sponge tool use in Indo-Pacific bottlenose dolphin matrilines, *Tursiops aduncus* [86]; and feeding on marine gastropod shells (“shelling”) in this same population of dolphins [87]. In other taxa, social learning has been documented in fish [88–90]; birds [20,91–96]; pollinators [97]; and primates [28,29,98–101]. All of these behaviours are performed individually, regardless of learning strategy or specific analytical test. It remains to be seen if and how the problems of using NBDA when the social network is largely defined by associations taking place during the behaviour of interest can be resolved, but the answer will be relevant to contexts beyond humpback whale cooperative bubble netting, such as to the well-documented occurrence of cooperative mud ring feeding in common bottlenose dolphins in Florida [41,42] and the Caribbean [102], and in Guiana dolphins, *Sotalia guianensis*, in the Cananéia estuary of São Paulo, Brazil [103]. One obvious way around this conundrum would be to use a social network constructed from observations outside the feeding context to predict the spread of the social behaviour, but in our case, this was not possible because while they are on a feeding ground the whales are doing little else.

Despite this caveat, even if individuals were to acquire the behaviour asocially, in order for them to ultimately feed cooperatively, there must be a phase of learning to cooperate. It seems highly likely that the development from a naïve individual, through to a fully cooperating bubble netter, involves some degree of social learning. Even if bubble netting is learned individually in a solo context, followed by homophily leading to involvement in the cooperative form, the division of tasks inherent in the cooperative form means that any transition from solo to cooperative bubble netting involves some learning about how and when tasks should be performed. This aspect of the learning must be social because it involves learning facilitated by the behaviour of others. Given that social learning can occur through interaction with the products of a behaviour, it seems feasible that a whale’s first exposure to cooperative bubble netting could involve learning from the product (the bubble net); but also by learning from the environment in which the behaviour occurs (a large enough prey patch to sustain a collective feed and the feeding calls emitted at particular frequencies) and from the coordinated movements of informed individuals. Their ability to exploit the nearshore habitat of this fjord system by bubble netting along the shoreline demonstrates a flexibility to respond to environmental, but perhaps also social, cues. This is particularly supported in the context that many individuals also perform other foraging strategies (e.g. flick feeding and lunge feeding) and some individuals switch between cooperative and solo bubble netting.

### Social cohesion

The interactions involved in cooperative bubble netting allow this feeding technique to serve as a mechanism for social cohesion. In a related study, the bubble netters of this same population were shown to have population-distinctive social behaviours [56]. Compared to the remainder of the population, known bubble netters were involved in longer-lasting and stronger social relationships, exhibited strong social preferences, were more socially connected, and exhibited greater centrality within the social network [56]. Bubble netters also exhibited higher rates of annual return and remained in the fjord system for longer durations in a given year [56]. These patterns underline that irrespective of the mechanism, social learning or homophily, the spreading bubble netting trait has a fundamental importance to the population’s social structure. Understanding the full effect of homophily on the diffusion of behaviours and the underlying social networks can be challenging, particularly because it can simultaneously accelerate the diffusion of a trait by helping to attain a critical mass, whilst also restricting the trait to homophilic groups within the network, depending on the strength of homophily at play [104]. Nonetheless, in addition to this example of humpback whales foraging in the KFS, it has been shown in another marine mammal species, populations of bottlenose dolphins in Laguna, Brazil and Shark Bay, Australia, that homophilic preferences tend to extend beyond the active performance of a behaviour to mediate the social network more generally [105,106].

### Solo bubble netting

The co-occurrence of solo and cooperative bubble netting is another example of behavioural plasticity in this population. Unlike the steady growth in numbers of informed individuals visible in Figure 2, the increase of new soloists (i.e. individuals observed solo bubble netting at least once) is slow until a sharp peak in the latter years of the study period (Fig. S2). This could be attributed to decreased prey availability or increased prey patchiness within the KFS in 2022 and 2023, given that we see the majority of these soloists switch back and forth between cooperatively and individually bubble netting (74% of whales known to solo bubble net have also been seen taking part in the cooperative form). A study relating prey abundance and distribution would therefore be an important next step.

### Ecological resilience

The North Pacific population of humpback whales had been steadily increasing at a relatively fast rate of 8% per year [69] until recent evidence of a drastic 20% decline in population size from 2012 to 2021 [67]. What sets humpback whales apart in their ability to rebound to pre-whaling numbers, which is not observed in cetacean species with whom they share a habitat [107– 110], might be explained by the interactions of their behavioural plasticity and their sociality [56], both underpinning rapid cultural evolution (or at times even ‘revolution’ as with the cultural transmission of song [47]). Since bubble netters are able to exploit schools of fish that naïve humpback whales cannot capture as effectively on their own [111], the diffusion of this behaviour has implications for this population’s ecological niche, carrying capacity, and resilience to environmental perturbations within Pacific Canada [56,112]. Individuals who have learned to bubble net are capable of exploiting distinct prey types (in the present case, euphausiids and herring) with highly disparate ecologies and phenologies, and do so both cooperatively and individually [55,107]. Such behavioural plasticity results in a resource base to be used more efficiently, which perhaps increases local carrying capacity. This, presumably, expands the viable foraging niche and can be the basis for further behavioural innovations.

In these ways, diffusion of techniques within social networks can bolster the recovery and resilience of a depleted population. Indeed, the dynamic nature of humpback culture may have played an important role in this species’ recovery from commercial whaling [69]. Cultural resilience may also play an important role in the future as humpback whales, like most cetaceans, face declines in habitat and population size [67,113–115]. Targeted efforts to protect a culturally distinct segment of a population may yield disproportionate benefits to diversity and resilience overall, but localised cultural loss can have an equally outsized effect [6]. In the KFS, a forecasted increase in large vessel traffic is expected to cause an increase in humpback whale ship-strike mortalities [116]. Should these mortalities affect core knowledge-holding, bubble netting individuals, a greater impact than expected may be seen on the seasonally resident population as a whole. For this reason, epicentres of distinct learned behaviours within management jurisdictions, such as the KFS in the Canadian Pacific, are strategic priority areas for conservation efforts [1,6]. To date, the consideration of cultural strains is rarely accounted for in conventional conservation frameworks [117– 119], but their importance is expected to grow as anthropogenic perturbations deteriorate marine ecosystems worldwide [113,114,120].

### Concluding remarks

We conclude that the spread of bubble netting in this population is intimately linked to, and potentially a causal factor in, its social network. We highlight issues around NBDA tests for social learning in a context of cooperation, as opposed to the social learning of behaviours performed alone. Furthermore, we show that the diffusion of bubble netting throughout northern British Columbia is not merely another example of this species’ predisposition for behavioural innovation; it is also a reminder that species recovery is more than a numerical process. As humpback whales undergo large fluctuations in population size in the northeastern Pacific [69,120], the distribution of behavioural knowledge will be an important factor in their recovery if it has been lost and is now being restored.

## ACKNOWLEDGEMENTS

We thank the Gitga’at First Nation for their stewardship and collaboration. We thank the Kitasoo/Xai’xais First Nations, the Haisla First Nation and the Heiltsuk First Nation for ongoing collaborative work. We thank Hermann Meuter, co-founder of Cetacea Lab. We thank Kevin Lala and Mauricio Cantor for insightful conversations that contributed to the interpretation of our results.

## Footnotes

1 Google Scholar and Web of Science search on the term “network-based diffusion analysis” was conducted on 17-10-2024. Study titles were screened for applications of NBDA and behaviours were examined on whether they were individual or group behaviours.

## BOX 1

**Positionality and reflexivity statement**

We comprise an international group of authors with a range of expertise in various biological disciplines who are collaborating on this project asking questions about sociality and culture in humpback whales foraging within the traditional territories of the Gitga’at First Nation, the Kitasoo and Xai’xais First Nations, the Gitxaała First Nation and the Haisla First Nation. We have an equal gender balance but come from a diverse set of backgrounds and perspectives. Authors NR and AD are Gitga’at members who have been dedicated to the study and protection of marine mammals in Gitga’at territory for approximately two decades. Author JW is the co-founder and CEO of the North Coast Cetacean Society (NCCS) and EMK, ÉOM, and GB are researchers with NCCS, which has been collaborating with the Gitga’at First Nation as an organisation since 2001, when permission to conduct research in the territory was granted and specific research agreements established. Authors EMK, ÉOM, GB, JW, LR and OEG recognise our intersecting social identities – we are all white and either living and working as settlers in Canada, or based at research institutions in Europe, from where we travel to spend time working with the Gitga’at in their unceded territory. We are striving to decolonise our research practices and acknowledge that we have much work to do on this still. Authors EMK, ÉOM, LR and OEG work in an academic system which affords us unearned privileges through its significant history of colonisation. All authors recognise that the biological sciences strive for objectivity, but we collectively believe it is critical in the process to decolonise ecology and evolution research that we address our inherent biases and belief systems. We are motivated by a deep care for Earth’s biodiversity, and in particular hope to see cetacean species co-existing with our own in perpetuity. We advocate for other researchers within ecology and evolution research to self-reflect on their own positionalities and relationships to land and community. We thank the Gitga’at Oceans and Lands Department and the Gitga’at community for their continued support and collaboration on whale research and conservation projects.

