## Supplementary Materials for "The diffusion of cooperative and solo bubble net feeding in Canadian Pacific humpback whales"

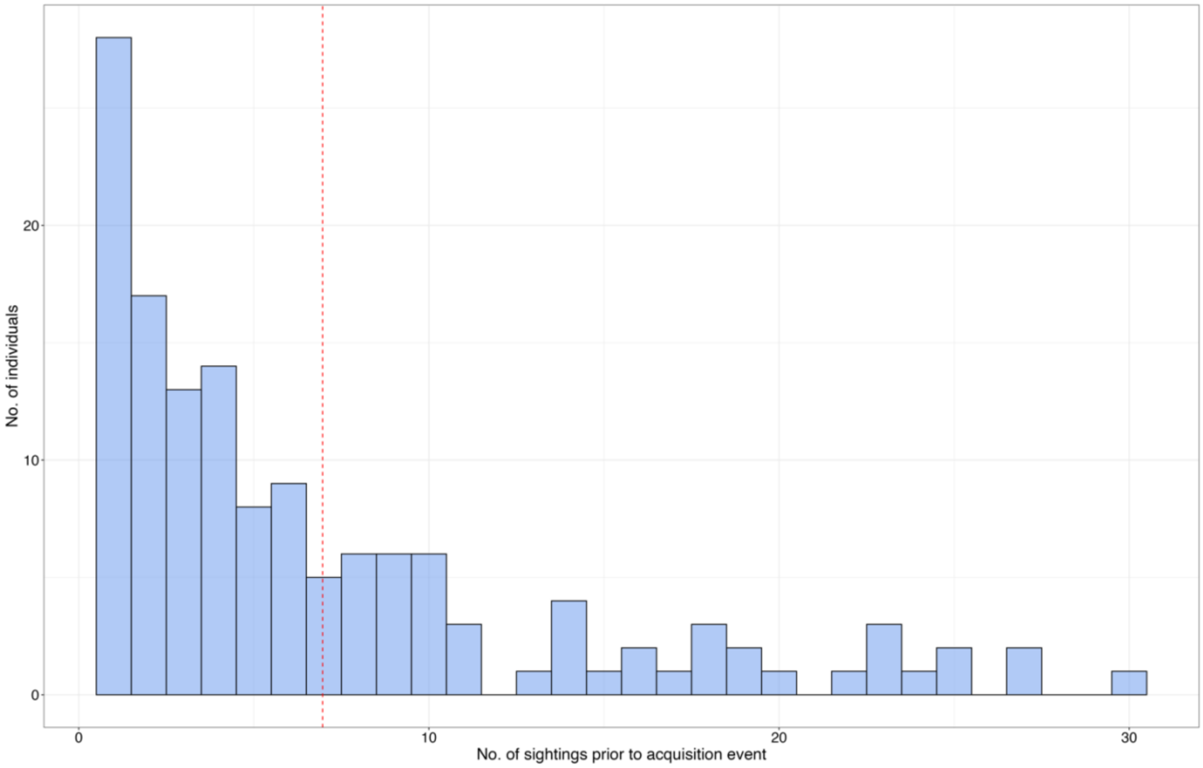


**Figure S1.** Distribution of the number of sightings of an individual prior to its first bubble net feeding event (i.e. its acquisition event), with data filtered to include only individuals seen at least 5 times in the study period. One outlier was removed with 67 sightings before it was observed bubble netting. Red dashed line denotes the mean ($\mu = 6.96$).

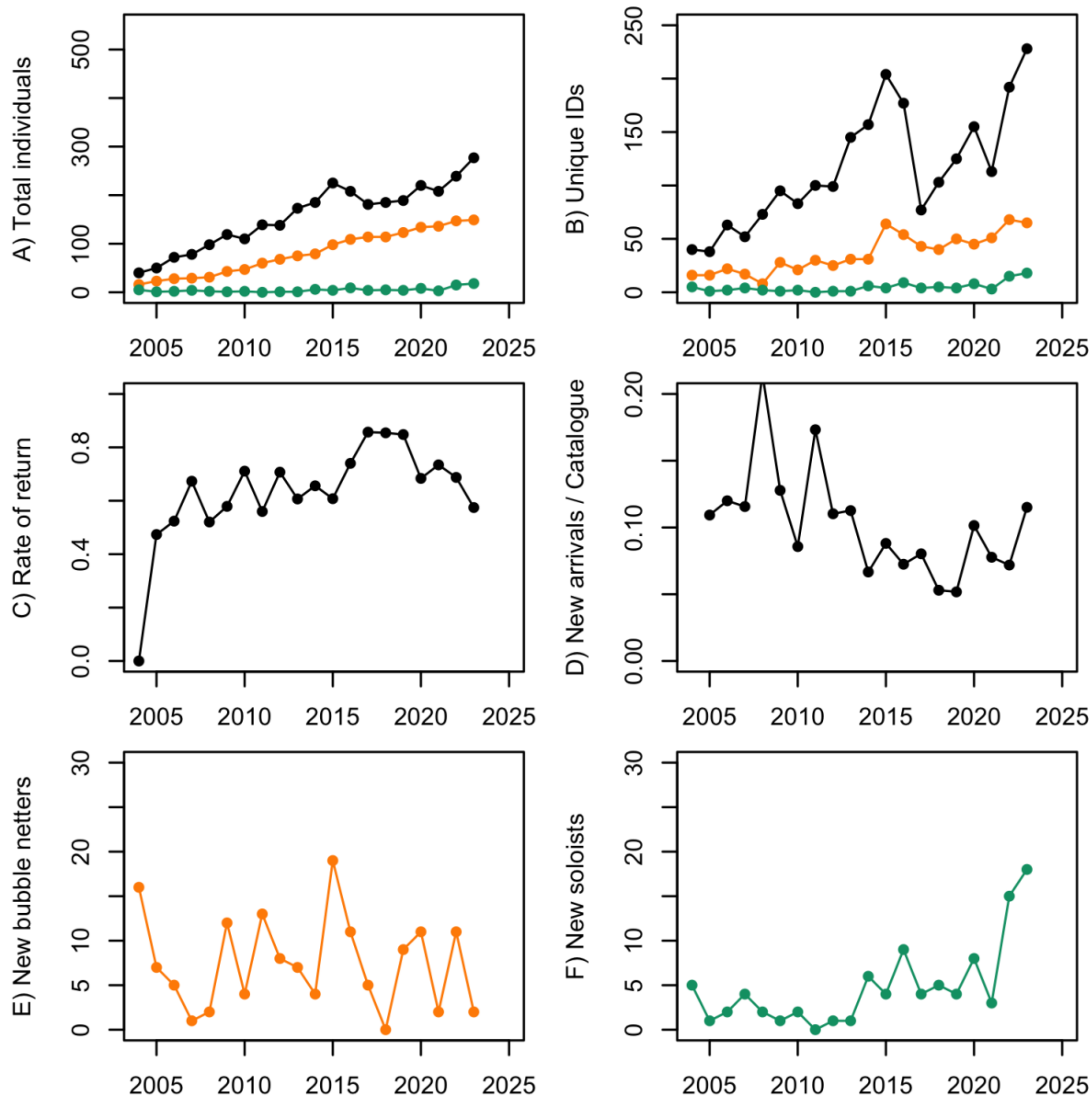


**Figure S2.** History of humpback whale observations from 2004 to 2023 for the entire studied population (black line), bubble netters (orange line) and solo bubble netters (green line). **A)** Discovery curve of newly identified individuals accumulating across the study period in black; while a diffusion curve is depicted in orange (and green for the diffusion of solo bubble netting). **B)** Uniquely identified individuals each year of the study period. The 2017 dip in unique IDs was likely due to reduced field effort in that year. **C)** The total rate of return of individuals seen prior in the study period. **D)** The proportion of the catalogue that are new arrivals each year. **E)** The number of new bubble netters each year. **F)** The number of new solo bubble netters ('soloists') each year.

*Additional information from outside of the study are*a

The hypothesis that bubble-net feeding is diffusing through the population, and potentially spreading south along the coast of British Columbia is supported by additional data collected in the central coast region near Bella Bella, where 118 individuals were identified in 141 encounters. The proportion of occasions in which feeding was observed that involved bubble net feeding was 0.12 in 2020, 0.25 in 2021 and 0.19 in 2022, with the number of individuals performing the bubble netting behaviour increasing from 5 to 10 to 15 in consecutive years. Only 60 out of 118 individuals (51%) identified were already known from the KFS catalogue. Of the 21 unique individuals observed bubble netting in the central coast, 6 had been previously observed performing this behaviour in the KFS, suggesting that it is not purely a movement of individuals that is responsible for these changes, but that perhaps the behaviour is being socially learned in the central coast from individuals that do transit between the two areas.

*Equation used to calculate the model-averaged parameter estimate as published in Eq. 6.12 of Burnham & Anderson (2002) [1].*

$$\widehat{\widehat{var}(\overline{\theta})} = \sum_{i=1}^{R} w_i \left[ \widehat{var}\left( \hat{\theta}_i \middle| g_i \right) + \left( \hat{\theta}_i - \hat{\overline{\theta}} \right)^2 \right]$$
